# Male-killing-associated bacteriophage WO identified from comparisons of *Wolbachia* endosymbionts of *Homona magnanima*

**DOI:** 10.1101/2022.06.12.495854

**Authors:** Hiroshi Arai, Hisashi Anbutsu, Yohei Nishikawa, Masato Kogawa, Kazuo Ishii, Masahito Hosokawa, Shiou-Ruei Lin, Masatoshi Ueda, Madoka Nakai, Yasuhisa Kunimi, Toshiyuki Harumoto, Daisuke Kageyama, Haruko Takeyama, Maki N. Inoue

## Abstract

The origin and mechanism of male-killing, an advantageous strategy employed by maternally transmitted symbionts such as *Wolbachia*, remain unclear. We compared genomes of four *Wolbachia* strains derived from *Homona magnanima*, a male-killing strain *w*Hm-t (1.5 Mb), and three non-male-killing strains, *w*Hm-a (1.1 Mb), *w*Hm-b (1.3 Mb), and *w*Hm-c (1.4 Mb). A *w*Hm-t-specific 76-kbp prophage region harboured two tandemly arrayed WO-mediated killing (*wmk*) gene homologs (*wmk-1*/*wmk-2* and *wmk-3*/*wmk-4*). Of these, *wmk-1* or *wmk-3* killed almost all *Drosophila melanogaster* individuals when transgenically overexpressed. Dual expression of *wmk-3* and *wmk-4* killed all males and rescued females. We propose a novel hypothesis wherein horizontally transmitted proto-*Wolbachia* with a single *wmk* killed both sexes, and tandem duplication of *wmk* allowed an evolutionary transition to a vertically transmitted symbiont, causing male-killing. Our study highlights the bacteriophage as a critical driver of the evolution of male-killing and argues for a conserved male-killing mechanism in diverse insects.

## Introduction

Insects harbour various endosymbiotic microbes, which occasionally interact with their hosts in mutualistic or parasitic manner (1–3). Male-killing (MK), the death of male offspring of infected female (4), is a reproductive manipulation caused by a diverse array of intracellular bacteria (5), microsporidia (6), and certain RNA viruses (7–8). MK is considered to have substantial ecological and evolutionary impact on insects as well as microbes, and the molecular mechanisms underlying MK have attracted extensive attention for decades (4, 9–12).

One of the best-studied MK endosymbionts is an alpha-proteobacterium *Wolbachia*, infecting approximately 40–60% of insects (13) and nematodes (14). MK *Wolbachia* strains have been identified from a variety of insect taxa which are not always phylogenetically closely related (15–16). Moreover, even closely related *Wolbachia* strains exhibit distinct phenotypes (17–20). *Wolbachia* is considered to have gained the ability to manipulate host reproduction multiple times by acquiring the bacteriophage WO (21–23). Indeed, the WO phage encodes cytoplasmic incompatibility (CI)-inducing factors (CifA and CifB), which result in the inability of sperm and eggs to form viable offspring (21-22, **Table 1**). Besides, the WO-mediated killing (*wmk*) gene, a putative transcriptional regulator located in the WO phage, is considered a candidate gene for MK because it induces a weak male lethality when transgenically overexpressed in *Drosophila melanogaster* (23). However, the *wmk* gene was derived from a non-MK *Wolbachia* (*w*Mel) naturally harboured by *D. melanogaster*, and the reason behind MK is not well understood. Moreover, two *wmk* homologs (i.e., one derived from *w*Suz, non-MK *Wolbachia* of *Drosophila suzukii*, and the other derived from *w*Rec, which does not induce MK in the native host *Drosophila recens* but causes MK when transferred into *Drosophila subquinaria* (24)) killed all males and females when transgenically overexpressed in *D. melanogaster* (25). These results suggested multiple host-dependent functions of *wmk.* Interestingly, female killing is mal-adaptive for maternally transmitted symbionts, and its evolutionary significance is not well understood. The WO phages frequently exhibit horizontal transfers among *Wolbachia* strains and are integrated into bacterial genomes as a temperate phage (prophage). Therefore, the WO phage may have been a critical driver of the evolution of *Wolbachia*-induced reproductive manipulations such as MK and CI.

**Table 1.** Reproductive phenotype-associated genes in Wolbachia and other male-killers

| Species and strain | Host insects | *Wolbachia*<br>phenotypes | Genes | Gene functions in transgenetically<br>overexpressed *Drosophila* | References |
| --- | --- | --- | --- | --- | --- |
| *Wolbachia pipientis*<br>wPip | *Culex pipiens* | CI*<sup>2</sup> | *cifA* and *cifB* | Recapitulated CI phenotype. | Beckmann et al. (21) |
| *Wolbachia pipientis*<br>wMel | *Drosophila melanogaster* | CI*<sup>2</sup> | *cifA* and *cifB*<br>*wmk* | Recapitulated CI phenotype.<br>Weak MK-like phenomenon (30% male death). | LePage et al. (22)<br>Perlmutter et al. (23) |
| *Wolbachia pipientis*<br>wRec | *Drosophila recens*<br>*Drosophila subquinaria* * <sup>1</sup> | CI*<sup>2</sup><br>MK*<sup>3</sup> | *wmk* | All death | Jaenike (24) and Perlmutter et al. (25) |
| *Wolbachia pipientis*<br>wSuz | *Drosophila suzukii* | CI*<sup>2</sup> | *wmk* | All death | Perlmutter et al. (25) |
| *Spiroplasma poulsonii*<br>(MSRO strain) | *Drosophila melanogaster* | MK*<sup>3</sup> | *spaid* | Recapitulated MK phenotype. | Harumoto and Lemaitre (26) |

\*1 wRec induces MK in the offspring of *D. recens* and *D. subquinaria*. Susceptibility to MK is recessive.
\*2 CI phenotype causes lethality in the offspring of *Wolbachia*-infected males and non-infected females.
\*3 MK phenotype causes lethality in males and leads to a female-biased sex ratio.

How microbes have acquired the ability to cause MK (origin) and how they exert the MK phenotype (mechanism) are key questions that remain unanswered (23–28). The bacterium *Spiroplasma poulsonii* (Mollicutes) causes MK by expressing a toxin-encoding *spaid* gene in *Drosophila melanogaster.* Nonetheless, *spaid* homologs have not been identified from other known male-killers, including *Wolbachia* (26). Besides, the expression of *Wolbachia*-induced MK depends on host species and *Wolbachia* genotypes (24, 29–31). For instance, the transinfection of the MK strain *w*Inn in *Drosophila innubila* does not cause MK in *D. melanogaster* and *D. simulans* (29). In contrast, a CI-inducing *w*CauA strain in *Ephestia* (=*Cadra*) *cautella* exhibits MK in *Ephestia kuehniella* (30). Since identical *Wolbachia* strains can cause different phenotypes depending on the host genetic background, comparative analysis of *Wolbachia* strains showing distinct phenotypes in a fixed host is an important approach to identifying MK gene(s).

Here, we focused on four *Wolbachia* strains, *w*Hm-t, *w*Hm-a, *w*Hm-b, and *w*Hm-c, harboured by a tea pest tortrix moth *Homona magnanima* (Tortricidae, Lepidoptera) (20, 32). *w*Hm-t (belonging to supergroup B) causes MK and is phylogenetically close to *w*Hm-c, which does not cause MK. On the other hand, *w*Hm-a and *w*Hm-b (both belonging to supergroup A) do not cause MK. Collectively, we explored the genetic characteristics of male-killers and the evolutionary processes in the acquisition of the MK phenotype.

## Results

### Comparative genomics of *Wolbachia* strains in *H. magnanima*

The MK-inducing *w*Hm-t was purified from the matriline W^T12^ by ultracentrifugation. Both PacBio RSII data (1.11 Gb, 110,635 reads, average read length: 7,603 bp) and Illumina data (1.44 Gb, 4,798,918 reads, paired-end 150 bp) were used to generate a polished 1,542,158 bp complete genome (**Table 2**). To construct genomes of *w*Hm-a, *w*Hm-b, and *w*Hm-c, an isofemale line derived from Yakushima Island (normal sex ratio for at least 30 generations, female ratio = 0.52, n = 1542) was subjected to single-cell analysis. Random sequencing of 384 libraries using single cells purified from males of the Yakushima matriline generated 6, 17, and 40 Illumina datasets for *w*Hm-a, *w*Hm-b, and *w*Hm-c, respectively. Following the quality assessment, GridION generated data of 138 Mb for *w*Hm-a (44,000 reads, average read length: 3,144 bp), 158 Mb for *w*Hm-b (48,000 reads, average read length: 3,281 bp), and 338 Mb for *w*Hm-c (128,000 reads, average read length: 2,637 bp) by using six single-cell amplicons for each strain. Genome assembling and polishing generated a complete genome of *w*Hm-c (1,447,668 bp) as well as draft genomes of *w*Hm-a (1,174,342 bp, 20 contigs) and *w*Hm-b (1,302,323 bp, 9 contigs) (**Table 2**).

**Table 2.** General characteristics of Wolbachia genomes

| Strain | wHm-a | wHm-b | wHm-c | wHm-t |
| --- | --- | --- | --- | --- |
| Supergroup | A | A | B | B |
| Phenotype | Not detected*<sup>1</sup> | CI*<sup>1</sup> | Shorten development time<br>Increase pupal weight*<sup>1</sup> | Male-killing*<sup>2</sup> |
| Contigs | 20 | 9 | 1 | 1 |
| Genome length (nt) | 1,174,342 | 1,302,323 | 1,447,668 | 1,542,158 |
| GC Content (%) | 35.28 | 35.12 | 34.01 | 34.06 |
| CDS | 1121 | 1,201 | 1,302 | 1,426 |
| Repeat region | Not identified | 200 | 128 | 177 |
| tRNA | 29 | 34 | 33 | 34 |
| rRNA | 3 | 3 | 3 | 3 |
| Completeness (%) | 85.3 | 98.0 | 99.7 | 100 |
| Contamination (%) | 3.1 | 2.7 | 0.3 | 0.3 |

\*<sup>1</sup> Based on Arai et al. (2019).
\*<sup>2</sup> Based on Arai et al. (2020).

High level of genome rearrangements were observed between *w*Hm-t, *w*Hm-a, and *w*Hm-b (< 50% homologies) (**Fig. 1A-B**). On the other hand, phylogenetically closely related *w*Hm-t and *w*Hm-c showed a high level of syntenic conservation (> 92% homology) in most parts of the genomes and were almost perfectly colinear with several inversions (**Fig. 1C**). *w*Hm-t and *w*Hm-c shared identical genes (84.9%, n = 1210) and homologs (> 90% identity, 94.9%, n = 1353) (**Extended Data Table 1**). Exclusively present in *w*Hm-t but not in *w*Hm-a, *w*Hm-b, and *w*Hm-c were six protein clusters constituting 14 proteins (**Extended Data Fig. 1**), which were, however, not possessed by MK *Wolbachia* found in other insects such as *Hypolimnas bolina* and *Drosophila* (i.e., *w*Bol1b (18), *w*Bor (23), and *w*Rec (23, 24)). On the other hand, two protein clusters that were absent in *w*Hm-t but present in *w*Hm-a, *w*Hm-b, and *w*Hm-c were hypothetical proteins and mobile elements. None of the putative proteins encoded by the MK Partiti-like virus in *H. magnanima* (7) or the MK protein Spaid encoded by the MK *Spiroplasma* in *Drosophila* (25) were homologous to any region of the *w*Hm-t genome.

**Fig. 1.**
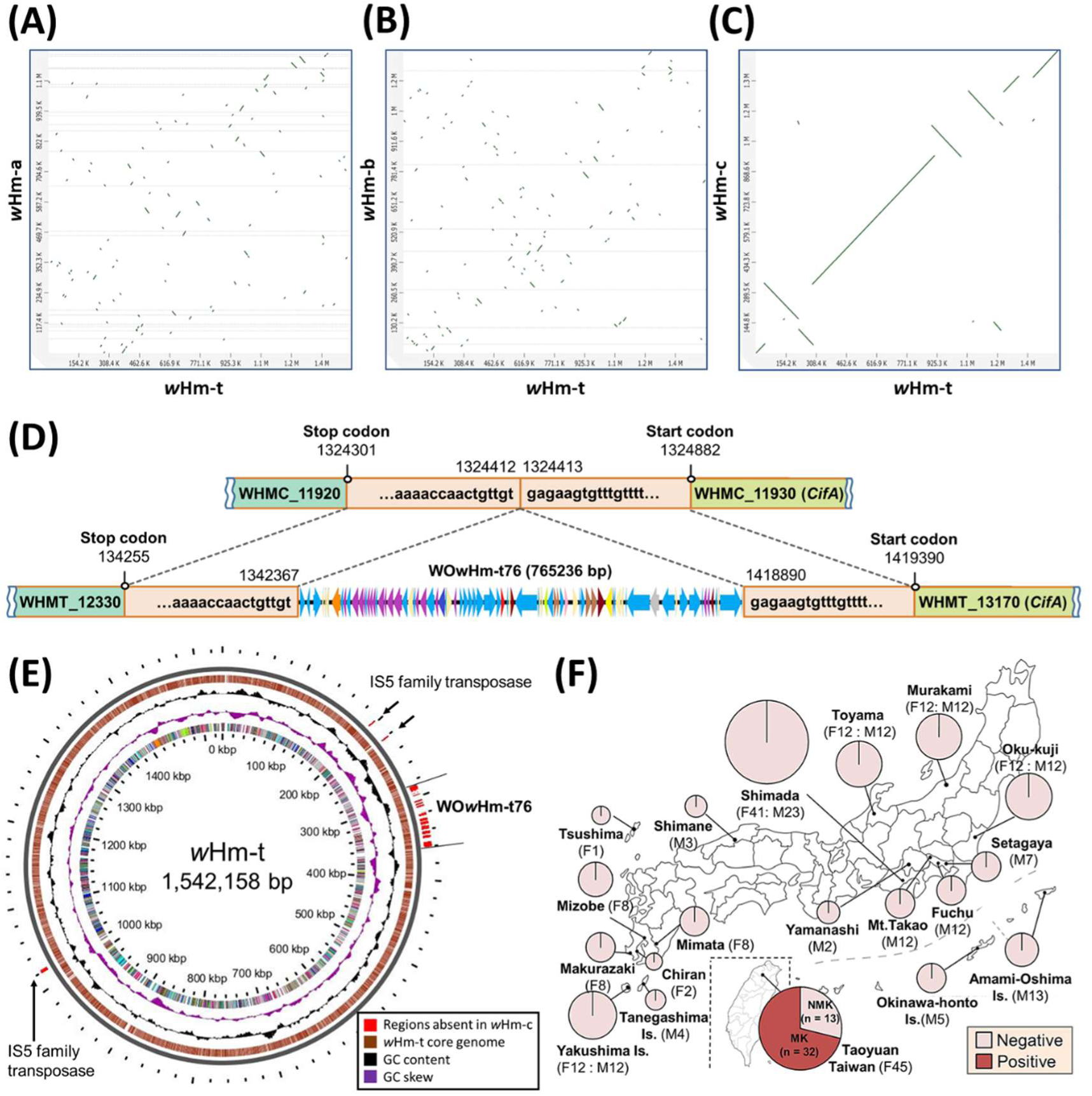
Genomic comparisons of *Wolbachia* strains in *H. magnanima*. Contigs of *w*Hm-a (A), *w*Hm-b (B), and *w*Hm-c (C) were compared and sorted based on the complete genome of *w*Hm-t. (D) An insertion, the WO*w*Hm-t76 region, in the *w*Hm-t genome. The wHmc_11920 of the *w*Hm-c encodes the IS110 family of transposase and has identical sequences of wHmt_12330 of the *w*Hm-t. wHmc_11930 of *w*Hm-c, encoding cytoplasmic incompatibility factor A (CifA), is identical to wHmt_13170 of *w*Hm-t. (E) Genomic loci, absent from *w*Hm-c genomes (n = 9) but conserved in *w*Hm-t genomes (n = 15), are highlighted in red on the *w*Hm-t genome (1,542,158 bp). Homologous genes or loci between *w*Hm-c and *w*Hm-t, such as the WO*w*Hm-t76 region, are not highlighted in red. (F) Detection of the WO*w*Hm-t76 region from *Wolbachia* infecting Japanese and Taiwanese *H. magnanima*. F: female individuals, M; male individuals.

### Identification of a *w*Hm-t-specific prophage region WO*w*Hm-t76

Notably, a 76-kb insertion (hereafter referred to as WO*w*Hm-t76), highly homologous to the *Wolbachia* phage WO (**Extended Data Figs. 2, 3**), was present in the *w*Hm-t genome (**Fig. 1D**) but not in *w*Hm-a, *w*Hm-b, and *w*Hm-c. Although several other prophage WO regions were identified on the *w*Hm-t and *w*Hm-c genomes (**Extended Data Figs. 2, 3**), further resequencing analysis confirmed that the WO*w*Hm-t76 region was conserved in *w*Hm-t (n = 15; Taiwan) but not in *w*Hm-c (n = 9; Japan). Other than the WO*w*Hm-t76 region, deleted or inserted loci in the *w*Hm-c were IS5 family transposases (**Fig. 1E**). To further confirm whether WO*w*Hm-t76 exists consistently in *w*Hm-t, the field-collected *H. magnanima* that were either infected with *w*Hm-c or *w*Hm-t were subjected to PCR that detected a sequence of the *cifB*-like gene (wHmt_13140) located in the WO*w*Hm-t76 region (**Fig. 1F**). All the Japanese *H. magnanima* derived from 18 populations consistently infected with the non-MK *w*Hm-c were negative (116 females and 129 males) for the *cifB*-like gene. On the other hand, all the 32 MK matrilines established from a single population of Taiwan were positive, and all the 13 non-MK matrilines established from the same population were negative for the *cifB*-like gene. In addition, *w*Hm-t typically showed higher titres than non-MK *w*Hm-c in *H. magnanima* (P = 0.003 by Steel–Dwass test; **Extended Data Fig. 4**), but *w*Hm-c did not cause MK even when its density was incidentally comparable to or higher than that of *w*Hm-t.

### WO*w*Hm-t76 harboured and expressed virulence-associated genes

The WO*w*Hm-t76 region was annotated to harbour 83 genes (wHmt_12340 to 13160), which contained *Wolbachia* titre/virulence-associated gene homologs such as ankyrin repeat-containing genes (n = 6, such as the *w*Hm-t-specific pk2 homolog wHmt_12420, **Extended Data Fig. 1**), virulence associated Octomom genes (n = 7), the *cifB*-like gene containing ankyrin repeats and papain-like peptidase domains (n = 1), and *wmk* homologs containing helix-turn-helix domains (n = 4) (**Fig. 2A**). RNA-seq and PCR assays confirmed that 51 genes in the WO*w*Hm-t76 region, including *cifB-like* and *wmk* genes, were expressed during embryogenesis and in the ovary of *H. magnanima* (**Extended Data Fig. 5**). Interestingly, the two tandemly arrayed *wmk* pairs in the WO*w*Hmt76 region (**Fig. 2B-C**) were conserved (100% nucleotide identity for each gene) and expressed in the MK *w*Bol1b (**Extended Data Fig. 5**). Besides, six *wmk* genes found in other *w*Hm-t genomic loci were identical to those of non-MK *w*Hm-c (100% nucleotide identity for each gene, **Fig. 2B**). Although *cifB* and *cifA* genes were tandemly arrayed in both CI-inducing *w*Mel and *w*Pip strains (21), only *cifB*-like gene (wHmt_13140, **Fig. 2D**) was found in the WO*w*Hm-t76 region. The Type IV *cifA* gene was conserved on their common region of the *w*Hm-c (wHmc_11930) and *w*Hm-t (wHmt_13170) strains. The CI-inducing *w*Hm-b encoded two tandemly arrayed Type I and Type II *cifA* and *cifB* gene pairs. Unexpectedly, *w*Hm-a, which does not induce CI, also possessed a tandemly arrayed Type I *cifA* and *cifB* gene pair.

**Fig. 2.**
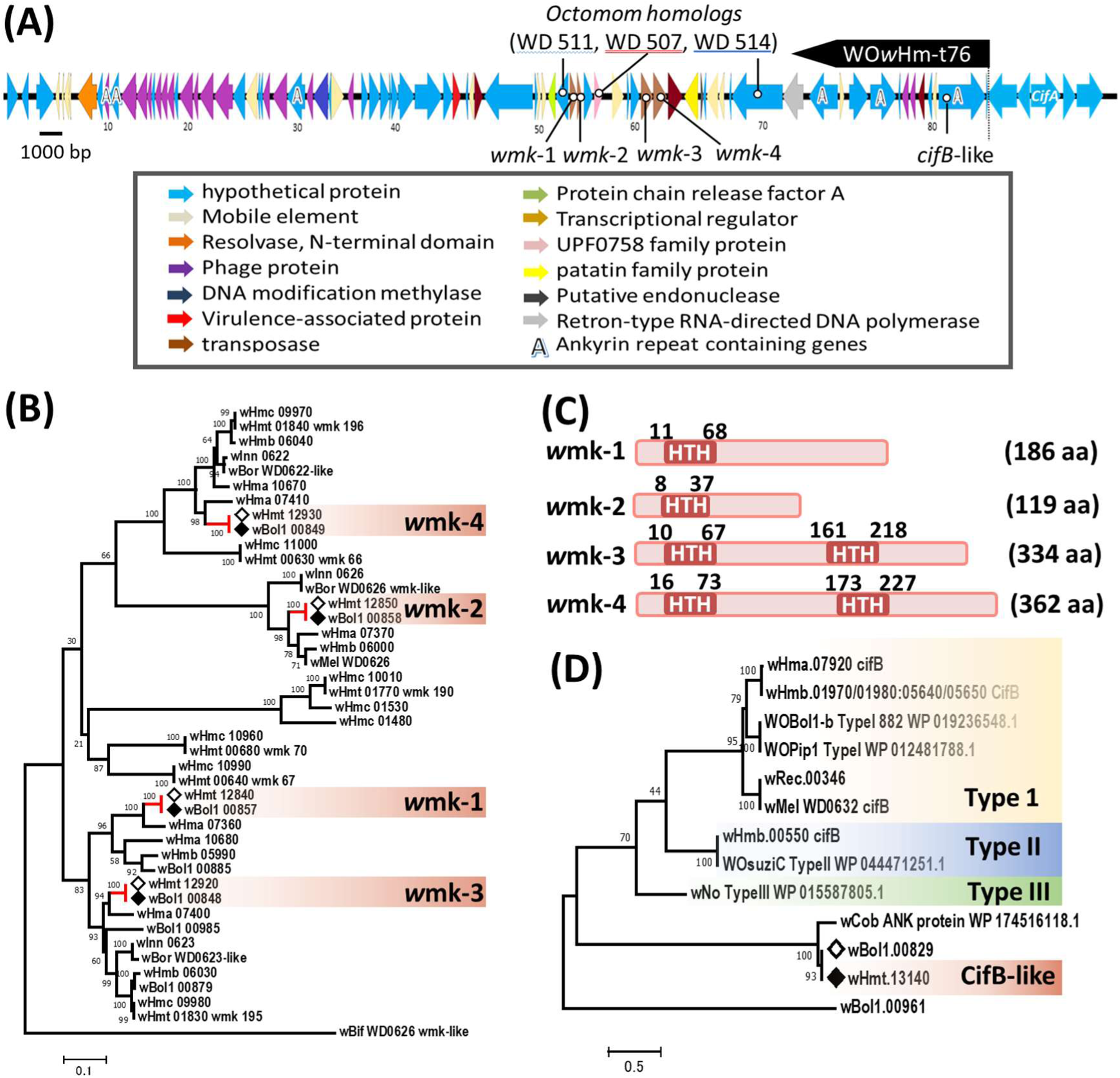
Phenotype-associated genes in the WO*w*Hm-t76 region. (A) Functions of genes in the WO*w*Hm-t76 region. (B) Phylogenetic trees of *wmk*. The *wmk* genes present in *w*Hm-t but absent in the *w*Hm-a, -b, and -c are highlighted with white diamonds (*wmk-1*, *wmk-2*, *wmk-3*, and *wmk-4*). Black diamonds indicate homologs of *wmk-1*, *wmk-2*, *wmk-3*, and *wmk-4* in the *w*Bol1b genome. (C) The domain structures of *wmk* annotated by InterPro. HTH: helix-turn-helix. (D) Phylogenetic trees of CifB homologs. Both *wmk* and *cifB* genes of *Wolbachia* strains were quoted from the NCBI database, and those of *w*Hm-a, -b, -c, and -t strains were manually annotated by local BLASTn and BLASTp in this study (Bit-score >100).

### Expression of some *wmk* genes singly killed both males and females in *Drosophila*

To assess whether the four *wmk* genes and a *cifB*-like gene on the WO*w*Hm-t-76 region affect insect viability, we used the *GAL4/UAS* system in *D. melanogaster*, as described by Harumoto and Lemaitre (26) and Perlmutter et al. (23), to zygotically overexpress each of the codon-optimised synthetic *wmk*-1 (561 bp), *wmk*-2 (361 bp), *wmk*-3 (897 bp), *wmk*-4 (1006 bp), and *cifB-like* (3558 bp) genes (**Fig. 3A**). When *wmk*-1 was overexpressed, only a small number of adult females (n = 52) and no males emerged in contrast to the control (982 females and 1014 males). When *wmk*-3 was overexpressed, almost no adults emerged (one female only) in comparison to the control (793 females and 888 males). Overexpression of *wmk*-2, *wmk*-4, and *cifB*-like genes did not affect survival or sex ratio compared to that in the controls (P = 0.89, 0.99, and 0.46 by Steel–Dwass test).

**Fig. 3.**
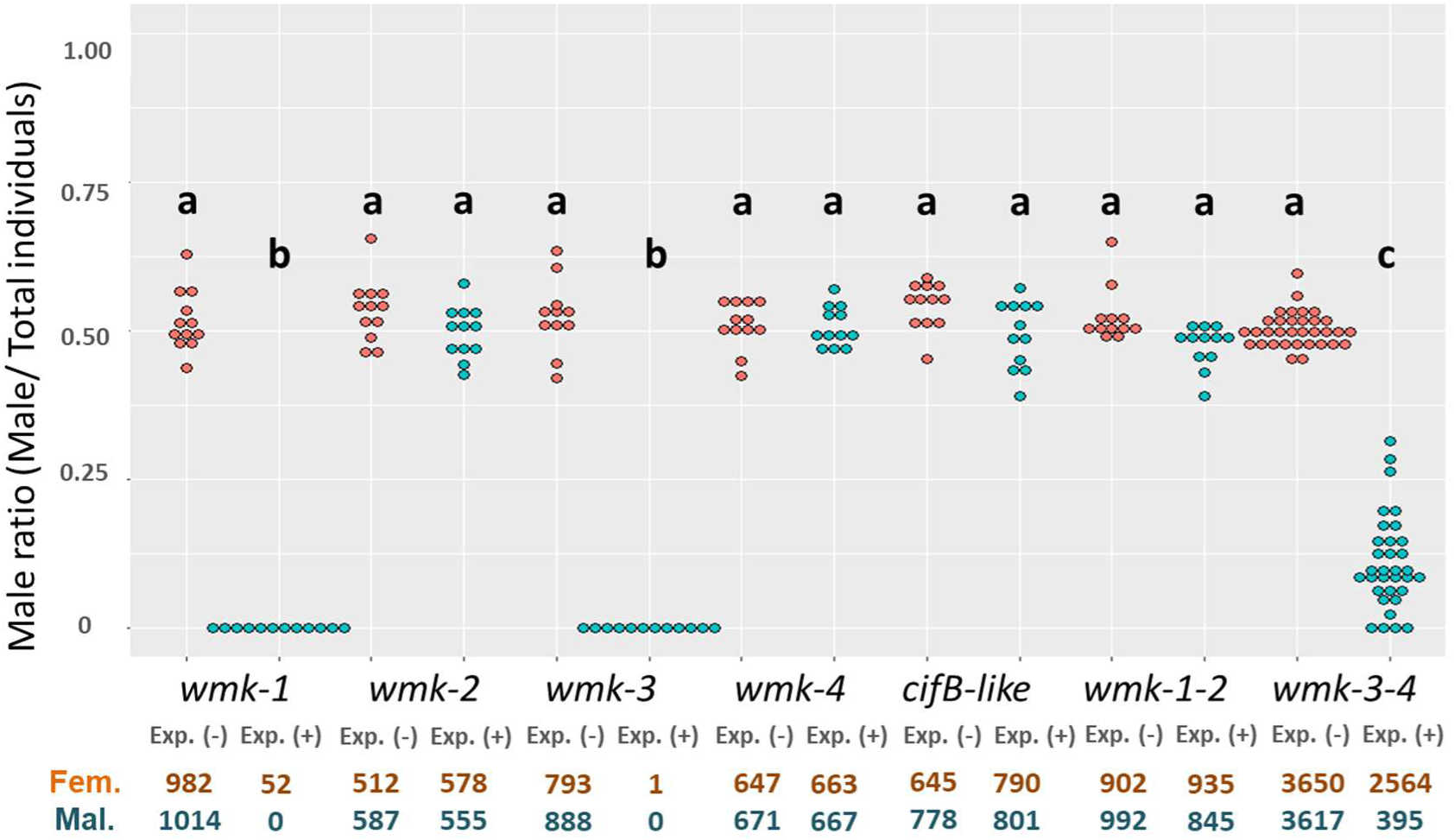
Expression of *wmk* genes caused lethality in *Drosophila melanogaster*. The male ratio of adult progeny obtained from crosses between the *actin–GAL4* line and seven *UAS* transgenic lines (*wmk*-1, *wmk*-2, *wmk*-3, *wmk*-4, *cifB*-like, *wmk*-1 and *wmk*-2, and *wmk*-3 and *wmk*-4; n = 12-32 independent crosses for each transgene). We counted the number of resultant offspring (females, orange; males, blue) having both *actin–GAL4* and *UAS* (+) and siblings having *UAS* and *Cyo (CyO)* as internal controls (−). The total number of adult counts for each genotype and sex are shown at the bottom. Different letters indicate statistically significant differences (Steel–Dwass test, P < 0.05). Dot plots show all data points individually. Exp.: expression; (-): non expressed; (+): expressed. Fem.: female progeny; Mal.: male progeny.

### Dual expression of *wmk* resulted in male-killing *in Drosophila*

To assess whether the dual expression of tandemly arrayed *wmk* pairs has additive or different effects on host lethality, we simultaneously overexpressed each pair of the *wmk* genes (i.e., *wmk*-1 and *wmk*-2 or *wmk*-3 and *wmk*-4) in *Drosophila*. Dual expression of *wmk-1* and *wmk-2* did not affect the viability and generated a normal sex ratio (male: 47.4%, female: 52.6%). Moreover, dual expression of tandemly arrayed *wmk-*3 and *wmk-* 4 increased the viability of female flies but led to low male ratios (male: 12.5%, female: 87.5%, **Fig. 3, Extended Data Fig. 6**).

## Discussion

In order to explore the origin and mechanism of MK in maternally transmitted symbionts, in this study, we compared the genomes of four *Wolbachia* strains derived from *H. magnanima* (a male-killing strain and three non-male-killing strains). We identified a 76-kb prophage region WO*w*Hm-t76, which is consistently present in the MK *Wolbachia* of *H. magnanima* but not in other *Wolbachia* strains that do not cause MK in a fixed host *H. magnanima*. Strikingly, a tandemly arrayed *wmk* gene pair encoded by WO*w*Hm-t76 caused strong MK when they were transgenically co-expressed in *Drosophila* flies.

The expression of MK is hypothesised to be mediated by factors, such as tropism or the expression level of genes present in MK *Wolbachia* (18). In *H. magnanima*, *w*Hm-t causes MK when infecting the host at a high density (20). In the current study, *w*Hm-t typically showed higher titres than non-MK *w*Hm-c in *H. magnanima*, but *w*Hm-c did not cause MK even when its density was incidentally comparable to or higher than that of *w*Hm-t. These results suggest that a high bacterial dose is not sufficient to induce MK in *H. magnanima*. *Octomom* genes, which enhance *Wolbachia* virulence and proliferation in *Drosophila* (33), on the *w*Hm-t-specific WO*w*Hm-t76 region may play a role in upregulating *w*Hm-t titre in *H. magnanima* to secure stable expression of MK. The WO phage possesses both a lytic and lysogenic nature and transfers genes laterally among *Wolbachia* strains (34–39); in the current study, the WO*w*Hm-t76 sequences were found in WO virions that were purified from the *w*Hm-t-infected host (**Extended Data Fig. 3**). We speculate that after the divergence of Japanese and Taiwanese *H. magnanima* populations, ancestral *w*Hm-c type *Wolbachia* in Taiwanese *H. magnanima* acquired the WO*w*Hm-t76 region through integration of a certain WO phage, which gave rise to the MK-inducing *w*Hm-t.

The *Wolbachia*-induced MK mechanism has remained unclear so far; however, we found that tandemly arrayed *wmk* genes caused strong male lethality with recovered female viabilities. The *Spiroplasma* endosymbiont in *D. melanogaster* recapitulates MK via the Spaid toxin that damages the male’s X chromosome using male-specific dosage compensation machinery, which normalises the expression of genes on the X chromosome between males (XY) and females (XX) (26). In contrast, *wmk-*1 and *wmk-*3 on the WO*w*Hm-t76 induced male as well as female lethality with different intensities in *Drosophila*; this is in line with the finding by Arai et al. (20) that *w*Hm-t also killed females when its titre was high in *H. magnanima*. The *wmk* gene (WD0626) derived from non-MK *w*Mel is reported to induce a weak MK in *Drosophila* (approximately 30% of males (23)). Moreover, *wmk* homologs of *w*Rec (MK in *D. subquinaria*) and *w*Suz (non-MK in *D. suzukii*), which only have several substitutions with WD0626, caused complete death (25). We speculate that the *wmk*-induced MK phenotype is represented by a different tolerance level between the sexes against *wmk* genes (**Fig. 4A**). Like the CI machinery of *Wolbachia*, the tandemly arrayed *wmk* genes might have combined effects, wherein one *wmk* shows toxicity toward both sexes and the other *wmk* acts as its suppressor. MK is considered to trigger diversification of the host sex determination pathways (10-11, 40-41). These systems in insects are diverse; for instance, the female-heterogametic system of Lepidoptera (including *H. magnanima*) and the male-heterogametic system of Diptera (including *D. melanogaster*) do not share any sex-determining genes known to date, except for the widely conserved *doublesex* gene. MK *Wolbachia* targets the dosage compensation system to damage male X chromosomes in *Drosophila* (42), whilst targeting the sex-determining gene cascade in *Ostrinia* and *H. magnanima*, thereby inducing improper dosage compensation in males (43–46). *Wmk,* assigned as a transcriptional regulator, may induce transcriptional effects leading to defects in host development, such as embryonic lethality (46) in Lepidoptera (*H. magnanima*) and Diptera (*Drosophila*) by a common but yet-unknown mechanism. The multiplication of *wmk* genes in *Wolbachia* genomes may have contributed to the diversification of sex-determining systems of arthropods.

**Fig. 4.**
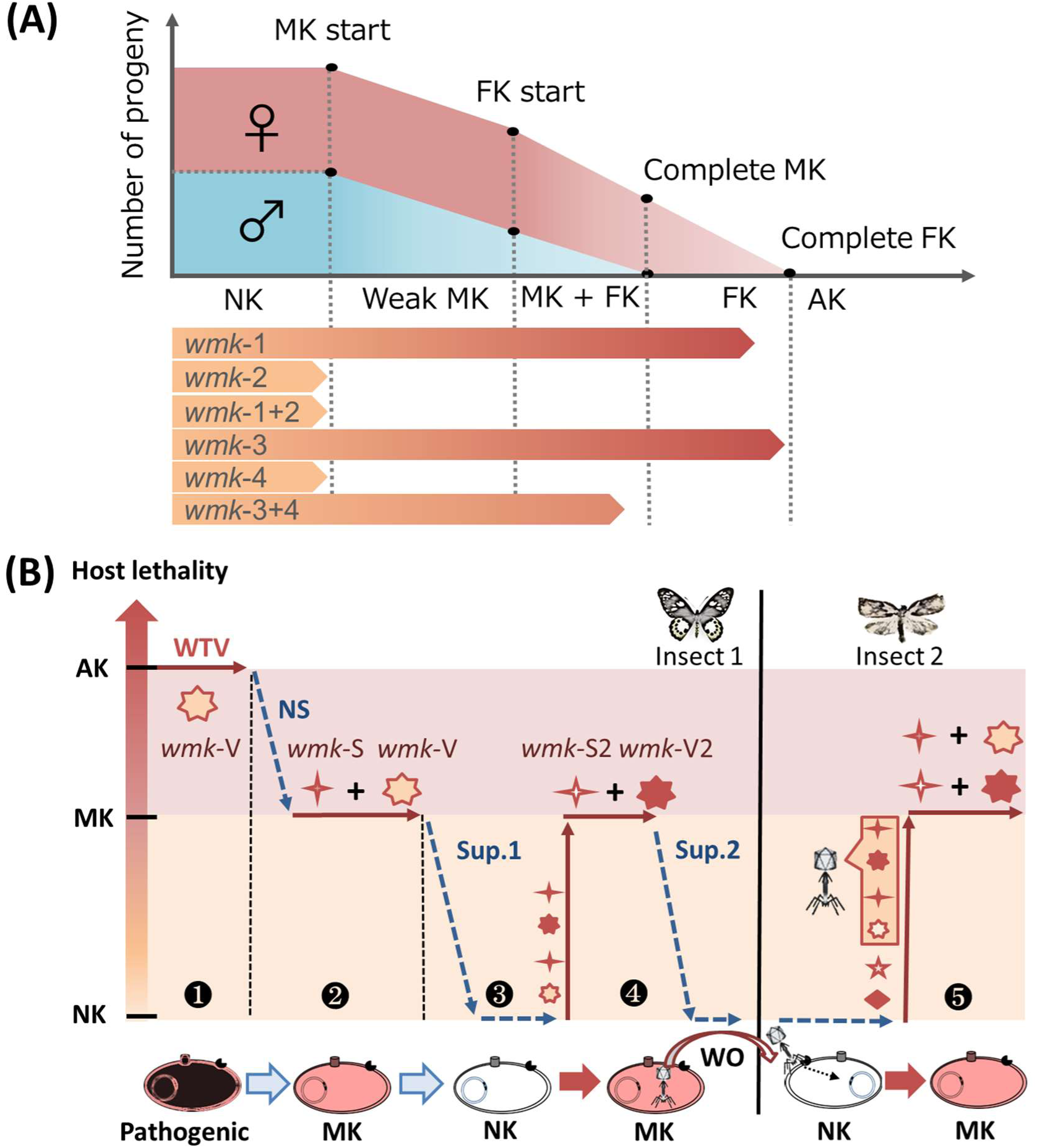
Models for *wmk*-induced lethality and evolutionary arms race between insects and *Wolbachia*. (A) A model of *wmk* inducing lethality. The *wmk*-1 and *wmk*-3 cause MK and female-killing (FK) during development. Males and females show differences in tolerance to virulent *wmk* genes such as *wmk*-1 and *wmk*-3, leading to differences in MK or FK start points. High expression of virulent *wmk* genes leads to nearly complete death (all killing, AK). The co-expression of tandemly arrayed virulent *wmk* (such as *wmk*-1 and *wmk*-3) and non-killing (NK) *wmk* (such as *wmk*-2 and *wmk*-4) leads to low virulence towards their hosts. The thresholds of killing phenotypes exerted by *wmk* genes could also be different due to host genotype or species. (B) Evolutionary arms race between insects and *Wolbachia* triggered by *wmk*-induced pathogenicity. Our current view of the *wmk*-induced pathogenicity system can be represented by a ‘zigzag’ model. In phase 1, *Wolbachia* deploys the virulent *wmk* gene (*wmk-V*) to induce *wmk*-triggered virulence (WTV) that contributes to an all-killing phenotype (AK). In phase 2, natural selection (NS) drives *Wolbachia* to reduce virulence toward females by shedding the *wmk-V* gene through acquiring an additional *wmk* gene (*wmk-S*) that suppresses a given *wmk-V* function, which leads to the MK phenotype of *Wolbachia*. In phase 3, insects develop suppressor (Sup.) towards the MK phenotype either by direct disablement of *wmk* functions or develop the host’s machinery to escape MK. In phase 4, duplications of *wmk* genes (*wmk-S2* and *wmk-V2*) result in MK, followed by the host’s new resistance to MK. In phase 5, the WO phage transmits *wmk* gene cassettes from a *Wolbachia* to a new strain, which leads to MK in a new host insect.

Why does maternally transmitted *Wolbachia* possess female lethal gene, which should bring about its own ruin? We propose a new hypothesis that a virulent *wmk* gene (e.g., *wmk-1* or *wmk-3*) was passed on by a horizontally transmitted pathogenic bacterium (i.e., ancestral *Wolbachia*) of insects. Rescue of females by the other *wmk* gene (e.g., *wmk-2* or *wmk-4*) should have allowed female offspring of the infected individuals to survive. This alteration *per se* can be regarded as the evolutionary transition of a horizontally transmitted pathogen to a vertically transmitted endosymbiont, which led to the prominent success of *Wolbachia,* as demonstrated by its current ubiquity. In general, virulent genes often exert duplications and substitutions through strong selective pressures (47–48). For the best-known reproductive phenotype CI, causative tandemly arrayed *cifB* and *cifA* genes are conserved among CI-inducing *Wolbachia* (21, 39). Strikingly, the present study demonstrated that non-CI-inducing *w*Hm-a strain also carried tandemly arrayed intact *cif* genes in its genome. These data suggest that the presence of *cif* genes cannot be an indicator of CI expression, despite the commonly held view that both *cifB* and *cifA* are necessary and sufficient to induce CI (21–22, 39). Likewise, the host-killing function of *wmk* genes may be widely conserved among diverse *Wolbachia* strains, including MK and non-MK strains. Indeed, MK *w*Hm-t and MK *w*Bol1b shared identical tandemly arrayed *wmk* gene pairs, whereas other MK (described in Perlmutter et al. (23)) and non-MK *Wolbachia* (e.g., *w*Hm-a, *w*Hm-b, and *w*Hm-c) strains generally possess multiple *wmk* gene cassettes regardless of their phenotype. It is tempting to speculate about an intense evolutionary arms race between *Wolbachia* and its host, which might have occurred in the past. An excessive female-biased sex ratio acts as a strong selection pressure to evolve host suppressors against MK in various insects (40). If MK were suppressed quantitatively, the production of a higher amount of MK toxins achieved by duplicating *wmk* gene cassettes would cancel the MK suppression (**Fig. 4B**). Subsequently, the development of host suppressors (10–11, 41) would trigger an evolutionary arms race between *Wolbachia* and hosts, which would increase the number of *wmk* genes in the *Wolbachia* genome. WO phages (such as WO*w*Hm-t76), vectoring *wmk* genes, may have contributed to the evolutionary arms race.

In summary, our study highlights the bacteriophage as a critical driver of the evolution of MK *Wolbachia*. Considering that *Wolbachia* has moved between different insect species and WO phages have moved between different *Wolbachia* strains (34–39), we speculate that the *wmk* genes in WO*w*Hm-t76 may induce MK in Lepidoptera (*H. magnanima*) as well as Diptera (*Drosophila*). However, we should be aware of the possibility that some other genes in WO*w*Hm-t76 do cause MK in *H. magnanima*. Further functional validations of these genes will contribute to an integrated understanding of the MK mechanisms induced by *Wolbachia* in diverse insects. Our discoveries offer insights into the molecular machinery of reproductive manipulations as well as evolutionary associations of MK *Wolbachia*, WO phages, and their host insects.

## Acknowledgments

We thank Professor Greg Hurst (Institute of Integrative Biology, University of Liverpool, Liverpool, UK) and Dr. Katsuhiko Ito (Tokyo University of Agriculture and Technology, Tokyo, Japan) for their kind advice. We also thank Hachiman-jyu Tea Factory (Yakushima, Kagoshima) for samplings of *H. magnanima* with the help of Mr. Hiroki Ishii and Ms. Mayu Nishino.

We wish to acknowledge support from the Japan Society for the Promotion of Science (JSPS) Research Fellowships for Young Scientists [Grant Number 19J13123 and 21J00895 to H. Arai], The Hakubi Project of Kyoto University (to T. Harumoto), JST ERATO [Grant Number JPMJER1902 to T. Harumoto], Nagase Science and Technology Foundation (to T. Harumoto) and MEXT KAKENHI Grant-in-Aid for Scientific Research(S) [Grant Number 17H06158] for H. Takeyama.

## Data Accessibility and Benefit-Sharing

### Data Accessibility

The sequence read data were deposited in DDBJ under accession numbers PRJDB13119 (BioProject) and SAMD00445397 to SAMD00445400 (BioSample). *Wolbachia* genomes (contigs) are available on the DDBJ database under the following accession numbers: *w*Hm-a (BQXF01000001 - BQXF01000020), *w*Hm-b (BQXG01000001 - BQXG01000009), *w*Hm-c (AP025639), and *w*Hm-t (AP025638). Homologs of *wmk*, *cif*, and Octomom genes (*w*Hm-a, *w*Hm-b, *w*Hm-c, and *w*Hm-t) and synthetic construct of *wmk* and *cif* genes were deposited in DDBJ under accession numbers LC701647 to LC701694.

### Benefits Generated

*H. magnanima* was collected from Tea Research and Extension Station (Taoyuan City, Taiwan) and imported with permission from the Ministry of Agriculture, Forestry and Fisheries (No. 27 - Yokohama Shokubou 891 and No. 297 - Yokohama Shokubou 1326). All collaborators are included as co-authors, and the results have been shared with the provider communities. More broadly, our group is committed to international scientific partnerships as well as institutional capacity building.

## Author Contributions

H. Arai conducted field surveys, genome sequencing, and data analysis; designed fly experiments; and wrote the original manuscript. H. Anbutsu coordinated the single-cell analysis and contributed to the entire discussion. Y. Nishikawa conducted the single-cell genome sequencing. M. Kogawa conducted the single-cell genome construction. K. Ishii helped with the genome construction. M. Hosokawa supported the entire single-cell analysis. SR. Lin organized the collection of insects in Taiwan and contributed to the discussion. M. Ueda collected Japanese *H. magnanima*. M. Nakai supported the entire experiments. Y. Kunimi sampled insects and contributed to the entire discussion. T. Harumoto designed and supported the fly experiments. D. Kageyama supported the fly experiments and revised the original manuscript. Lastly, H. Arai, H. Takeyama, and M.N. Inoue took responsibility for the decision to submit the manuscript for publication and managed the experiments and discussion.

## Materials and Methods

### Sampling of *H. magnanima*

To construct *Wolbachia* genomes, we used two *H. magnanima* lines: a MK line (W^T12^), infected with *w*Hm-t, and a normal line (Yakushima), triply infected with *w*Hm-a, *w*Hm-b, and *w*Hm-c. For resequencing of *Wolbachia*, we used field-collected Japanese and Taiwanese *H. magnanima* (**Extended Data Table 2**).

### Genome construction of *Wolbachia*

For sequencing of the *w*Hm-t genome, *Wolbachia* cells purified from the W^T12^ line were used for DNA extraction following Duplouy et al. (18) and Iturbe-Ormaetxe et al. (49). Extracted DNA was whole-genome-amplified (WGA) with the REPLI-g Mini Kit (Qiagen) following the manufacturer’s protocol and was sequenced using PacBio RSII and Illumina (150 bp paired-end) platforms. PacBio RSII data were assembled using Canu (50), and subsequent polishing using Illumina data with minimap2 (51) and Pilon (52) generated the complete genome of *w*Hm-t. For genome sequencing of *w*Hm-a, *w*Hm-b, and *w*Hm-c, *Wolbachia* cells purified from the Yakushima line were subjected to single-cell analysis using MiSeq (75 bp paired-end) and GridION, as described by Chijiiwa et al. (53) and Nishikawa et al. (54). The sequence data from GridION were assembled by Canu (50) and were polished via Pilon (52) using MiSeq data. *Wolbachia* genomes were annotated via DFAST (55). Phage WO infections were annotated with Phaster (56). *Wolbachia wmk*, *cifA,* and *cifB* genes were annotated using BLAST searches and utilized to construct phylogenetic trees using MEGA7 (57) by using maximum likelihood with bootstrap re-sampling of 1,000 replicates. Homology between the *spaid* (26) versus *w*Hm-t genes or genes of the Partiti-like viruses, OGV (7), versus *w*Hm-t genes were analysed by BLAST searches.

### Resequencing analysis and detection of the WO*w*Hm-t76 region

DNA extracted from *H. magnanima* harbouring *w*Hm-t (n = 15, Taiwan) and *w*Hm-c (n = 9, Japan) was subjected to resequencing on an Illumina platform (150 bp paired-end). Resequenced data, mapped to the *w*Hm-t genome (W^T12^) using minimap2 (51), were converted to consensus genomes using SAMtools (58), followed by synteny analysis using GView (https://server.gview.ca). Homologies of the WO*w*Hm-t76 region and other WO phages were visualized using Easyfig (59) and Mauve (60). The presence or absence of the WO*w*Hm-t76 region in field-collected *H. magnanima* was assessed by PCR amplifying a *cifB-*like gene (**Extended Data Table 3**) in the WO*w*Hm-t76 region with the Emerald Amp Max Master mix (TaKaRa) at 94°C for 3 min, 35 cycles of 94°C for 30 sec, 62°C for 30 sec, 72°C for 3 min, and final extension at 72°C for 7 min.

### WO purification, observation, and phylogenetic analysis

WO virions were purified as described by Fujita et al. (7). The recovered virions were dialyzed using PBS and were loaded on membrane-coated transmission electron microscopy grids (Nissin EM) and stained with 2% uranyl acetate solution (pH 7.0). The samples were observed using a JEM1400 Plus (Joel). DNA extracted from purified virions was subjected to PCR detection for genes of WO (*cifB*-like and minor capsid protein [ORF7]), *Wolbachia* (*wsp*), and *Homona* (*β-actin*). Phylogenetic trees of WO minor capsid protein genes of *Wolbachia* strains were estimated using MEGA7 (57) by using maximum likelihood with bootstrap re-sampling of 1,000 replicates.

### Gene expression of *w*Hm-t

Gene expression dynamics of *w*Hm-t in *H. magnanima* were assessed by RNA-seq and RT-PCR using RNA extracted from W^T12^ adults or egg masses of 12, 36, 60, 84, and 108 hours post oviposition, as described by Arai et al. (61). Gene expression levels were assessed using minimap2 (51) and iDep9 (http://bioinformatics.sdstate.edu/idep/). Specific primers for *wmk* (*wmk-1*, *wmk-2*, *wmk-3*, and *wmk-4*) and *CifB* genes (**Extended Data Table 3**) were used for RT-PCR. Expression levels of the *wmk* genes were further tested using a female *Hypolimnas bolina* (collected from Ishigaki Is., Okinawa) harbouring *w*Bol1b.

### Fly stocks and expression experiments

The *wmk* and *cifB*-like genes in the WO*w*Hm-t76 region were synthesised with codon optimisation for *D. melanogaster.* To express two tandemly arrayed *wmk* pairs, either (i) *wmk*-1 and *wmk*-2 or (ii) *wmk*-3 and *wmk*-4 were conjugated using a T2A peptide as described by Beckmann et al. (22). Transgenic fly lines were established using the standard microinjection method for phiC31 integrase transformation into the third chromosome at BestGene Inc. (Chino Hills, CA, UAS) using the synthetic constructs of these genes in the pUASZ 1.1 plasmid. To transgenically overexpress the *wmk* and *cifB*-like genes, *D. melanogaster* line *actin5C–GAL4* [*actin–GAL4*; BDSC 4414], cultured in a tetracycline-containing medium (0.05% [w/v]) for one generation to eliminate native *Wolbachia,* were crossed to homozygous *UAS* transgenic flies harbouring *wmk* or *cifB*-like genes. The numbers of offspring expressing transgenic genes (*actin-GAL4/UAS*) and non-expressing genes (*Cyo/UAS*) were all counted and were compared using Steel–Dwass test in R software v4.0 (https://www.r-project.org/).

---

**Extended Data Fig. 1.**
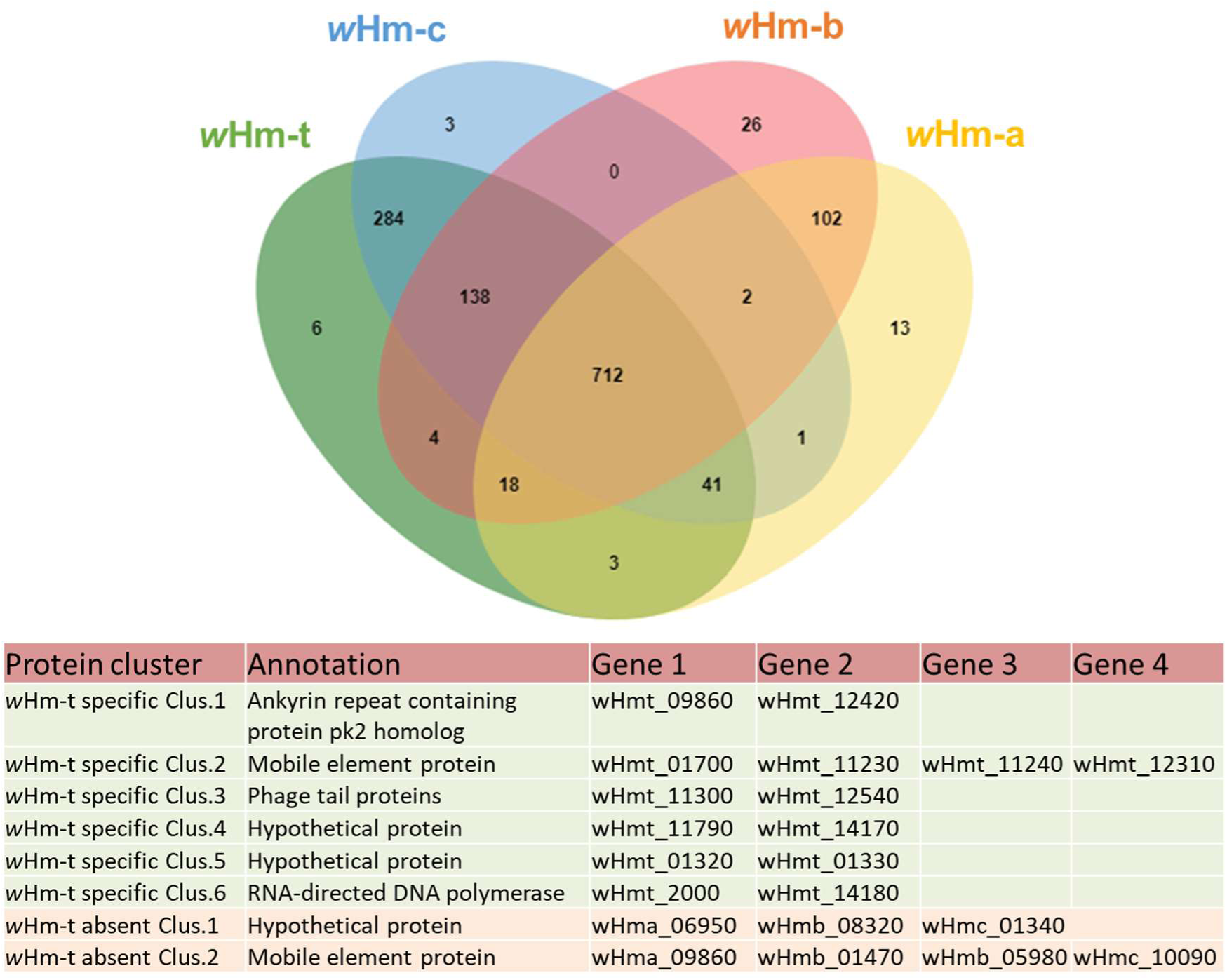
Genetic comparisons of four *Wolbachia* strains in *H. magnanima*. Venn diagram of protein clusters of the *w*Hm-a, *w*Hm-b, *w*Hm-c, and *w*Hm-t strains. Numbers indicate gene clusters. Descriptions of *w*Hm-t-specific protein clusters (n = 6) and protein clusters specifically absent in the *w*Hm-t genome (n = 2) are highlighted in the table shown below the diagram.

**Extended Data Fig. 2.**
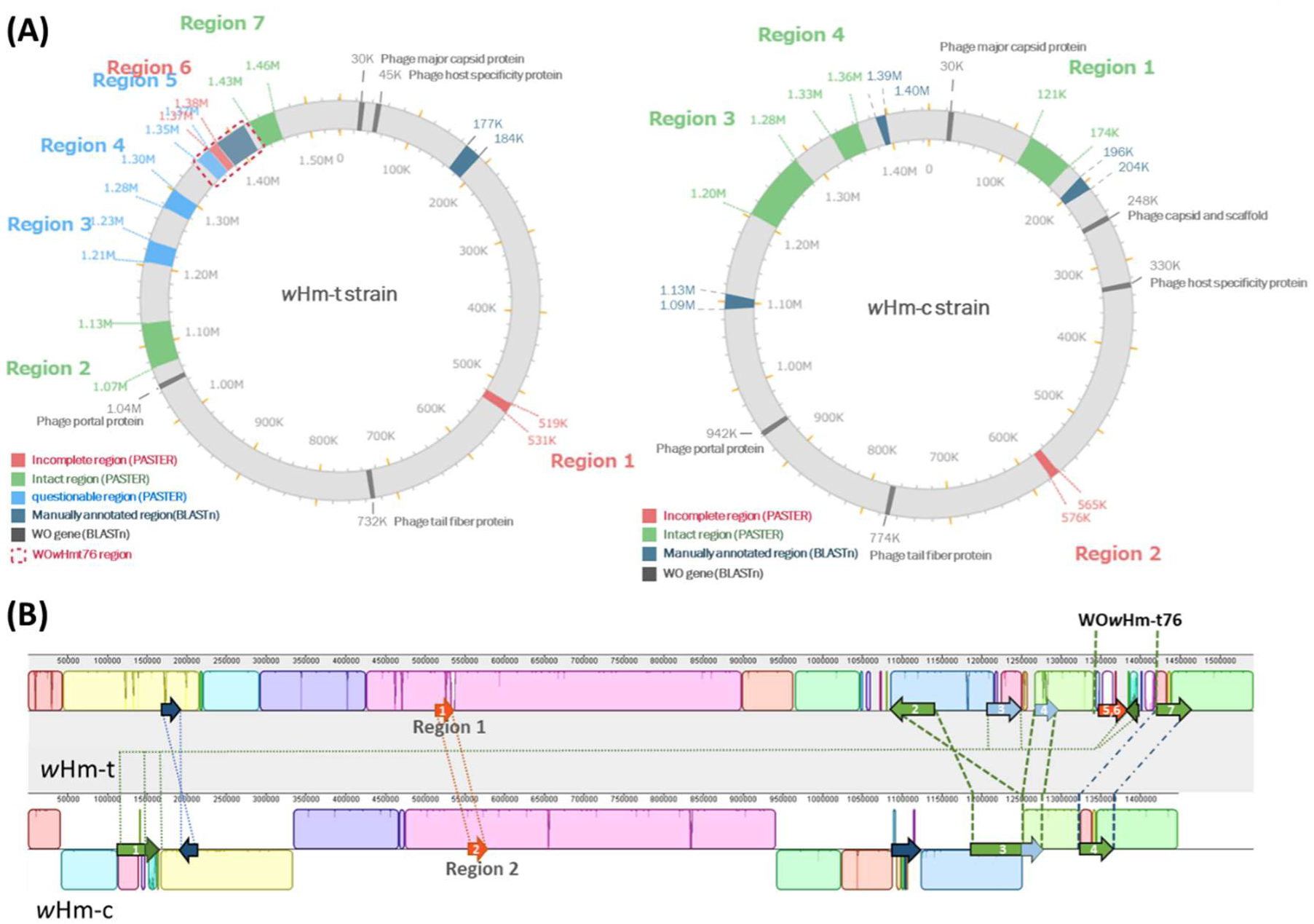
Comparisons of WO in *w*Hm-t and *w*Hm-c. (A) WO prophage regions were highlighted on both *w*Hm-t and *w*Hm-c genomes by PHASTER and manual BLASTn searches. (B) Homologies of WO regions on the *w*Hm-c and *w*Hm-t genomes. Blocks shown in the same colours indicate homologous genomic loci. The arrows indicate the WO region. Homologies are indicated with broken lines.

**Extended Data Fig. 3.**
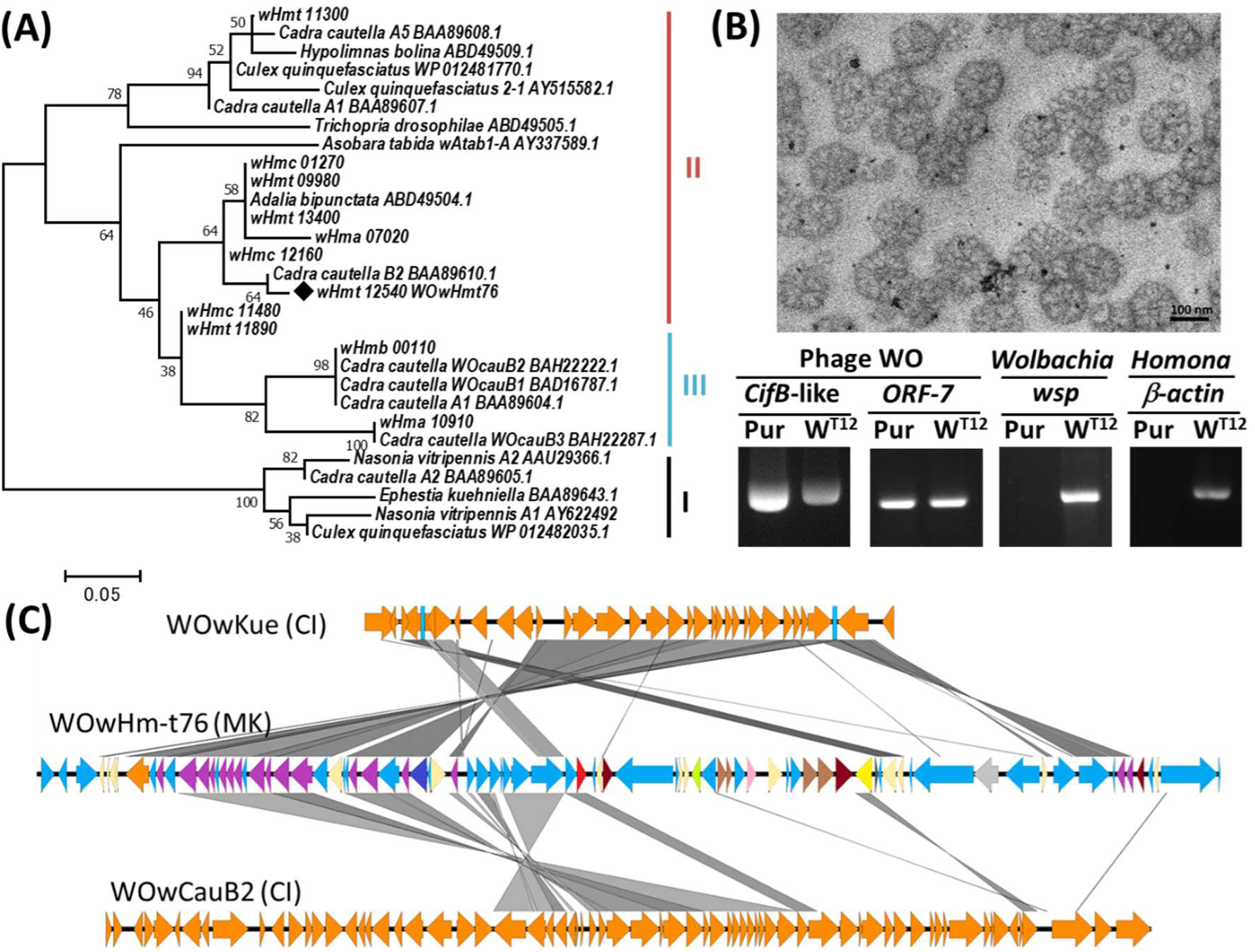
Phylogeny and morphology of WO. (A) Phylogeny of *Wolbachia* phage WO, based on the minor capsid protein (ORF7). A maximum-likelihood tree (1,000 bootstraps) inferred from amino acid sites of *Wolbachia* strains is shown. Sequence accession numbers are indicated in the figure. I, II, and III represent the ORF7 clusters identified by Bordenstein and Wernegreen (37). (B) Purified WO phage particles visualized using a transmission electron microscope (TEM) with uranyl acetate staining. WO genes (ORF-7 and *cifB*-like) were detected from DNA extracted from purified WO particles (Pur) and *H. magnanima* harboring *w*Hm-t (W^T12^). Both *Wolbachia* (*wsp*) and host (*b-actin*) were negative for the fraction. (C) Homologies of WO*w*Hmt-t76 and two *Wolbachia* phage WOs. The WOs identified from *w*Kue (AB036666) and *w*CauB (AB478515) were compared by BLASTn.

**Extended Data Fig. 4.**
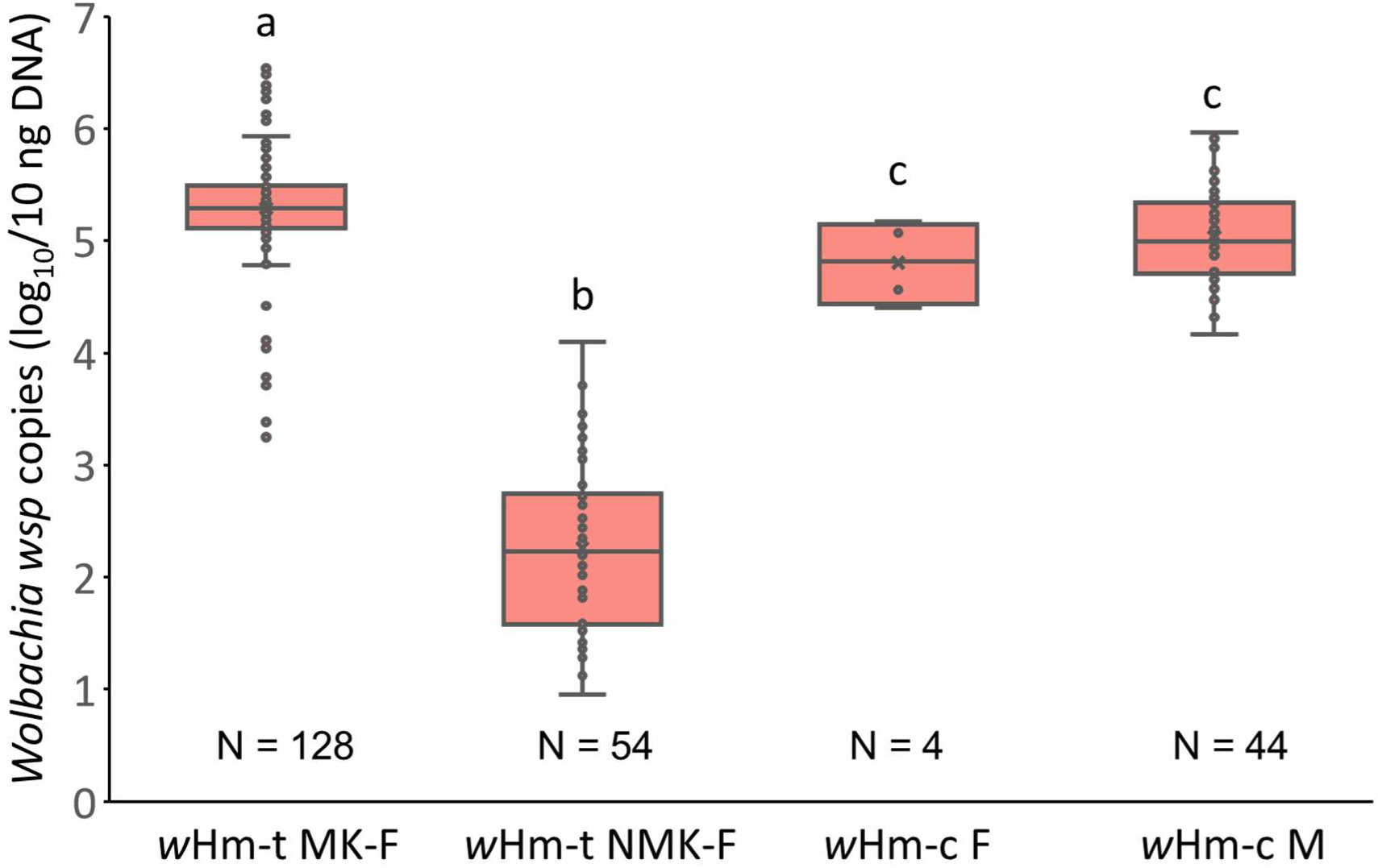
Titres of *Wolbachia* in Taiwanese *w*Hm-t-infected and Japanese *w*Hm-c-infected host individuals. *Wolbachia* copies (*wsp*) in *w*Hm-t-infected females of Taiwanese host lines (collected by Arai et al., 2021) and *w*Hm-c-infected Japanese populations surveyed in this study were compared. The centreline within the box represents the median. The upper and lower boundaries of the box indicate the upper quartile and lower quartile, respectively. The sample size (number of individuals) in each data point is displayed below the plots. Different letters indicate significant differences between groups (Steel-Dwass test, *p* < 0.05). MK: male-killing, NMK: non-male-killing, F: female. M: male.

**Extended Data Fig. 5.**
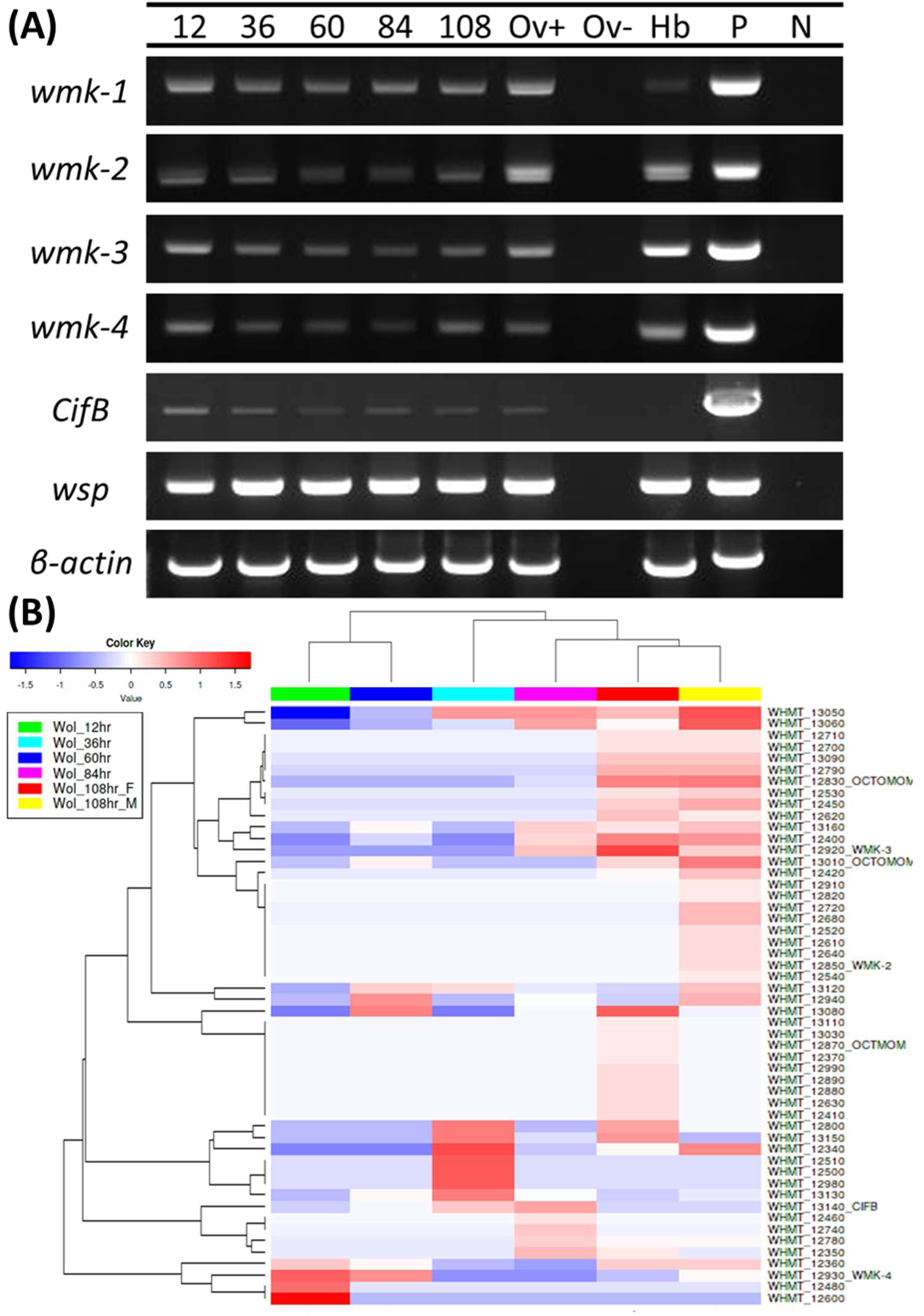
Expression of genes in the WO*w*Hmt-t76 region. (A) RT-PCR detection for the *wmk* and *cifB*-like genes. 12, 36, 60, 84, and 108 indicate hours post oviposition of *H. magnanima* egg masses. Ov+: female ovary of *H. magnanima*; Ov-: non-reverse-transcribed samples (RT minus, control) using RNA extracted from the female ovary of *H. magnanima*; Hb: female ovary of *Hypolimnas bolina*; P: DNA extracted from a W^T12^ adult female (positive control); N: negative control (water). (b) Expression of genes on the WO*w*Hmt-t76 region throughout embryogenesis. Mapping counts of each gene obtained by RNA-seq data (*H. magnanima* egg masses 12–84 hours post oviposition and 108-hour male or female embryos) were analysed. Expression levels are highlighted in blue (low) and red (high). F: female; M: male.

**Extended Data Fig. 6.**
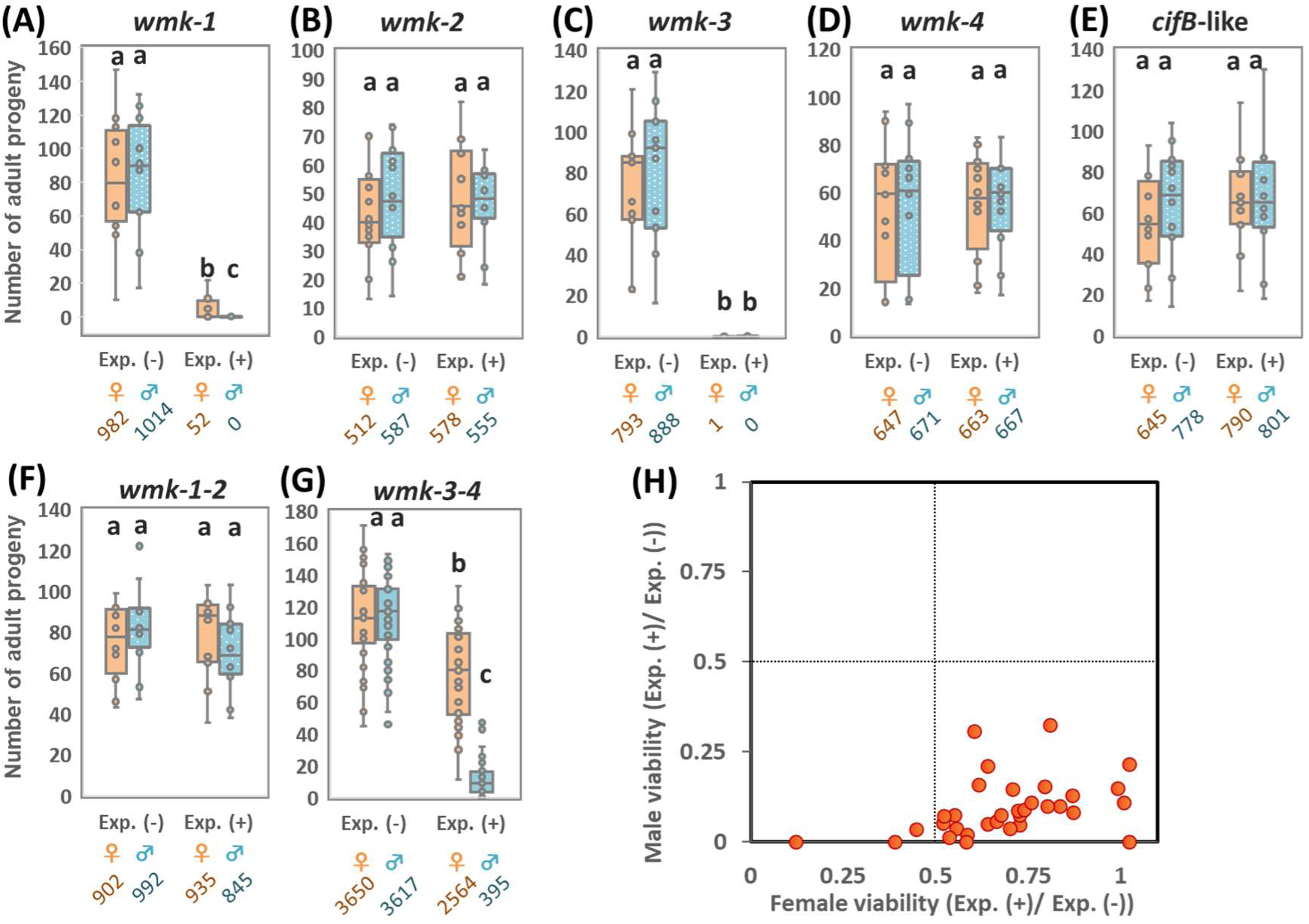
Number of female and male progenies in *D. melanogaster* expressing genes and their correlation. The number of adult progenies obtained from crosses between the *actin–GAL4* line and seven UAS transgenic lines: *wmk*-1 (A), *wmk*-2 (B), *wmk*-3 (C), *wmk*-4 (D), *cifB*-like (E), *wmk*-1 and *wmk*-2(F), and *wmk*-3 and *wmk*-4 (G) (n = 12-32 independent crosses for each transgene). We counted the number of resultant offspring (females, orange; males, blue) having both *actin–GAL4* and *UAS* (+) and siblings having *UAS* and *Cyo* as internal controls (−). The total number of adult counts for each genotype and sex are shown at the bottom. Exp.: expression; (-): non expressed; (+): expressed. (H) Dot plots of male and female viabilities in each independent cross for *wmk*-3 and *wmk*-4 dual expression.

**Extended Data Table 1.** Homologies of total 1,426 wHm-t genes versus genes of other non-MK strains in *H. magnanima*

| Homologies *<sup>1</sup> | *wHm-a* | *wHm-b* | *wHm-c* |
| --- | --- | --- | --- |
| Genes with 100% identity | 11 | 0 | 1,210 |
| Genes with $\geq 95\%$ identity | 64 | 39 | 1,328 |
| Genes with $\geq 90\%$ identity | 253 | 214 | 1,353 |
| Genes with $\geq 80\%$ identity | 835 | 794 | 1,377 |
| Genes with $\geq 70\%$ identity | 858 | 820 | 1,379 |
| Genes with no homologies*<sup>2</sup> | 568 (387) | 606 (298) | 47 (12) |

\*<sup>1</sup> The identity of genes was analysed by using BLASTn.
\*<sup>2</sup> Numbers to the left of the parentheses indicate genes specific for *wHm-t*, and numbers shown in parentheses indicate the genes specific for the compared non-MK strain.

**Extended Data Table 2.**
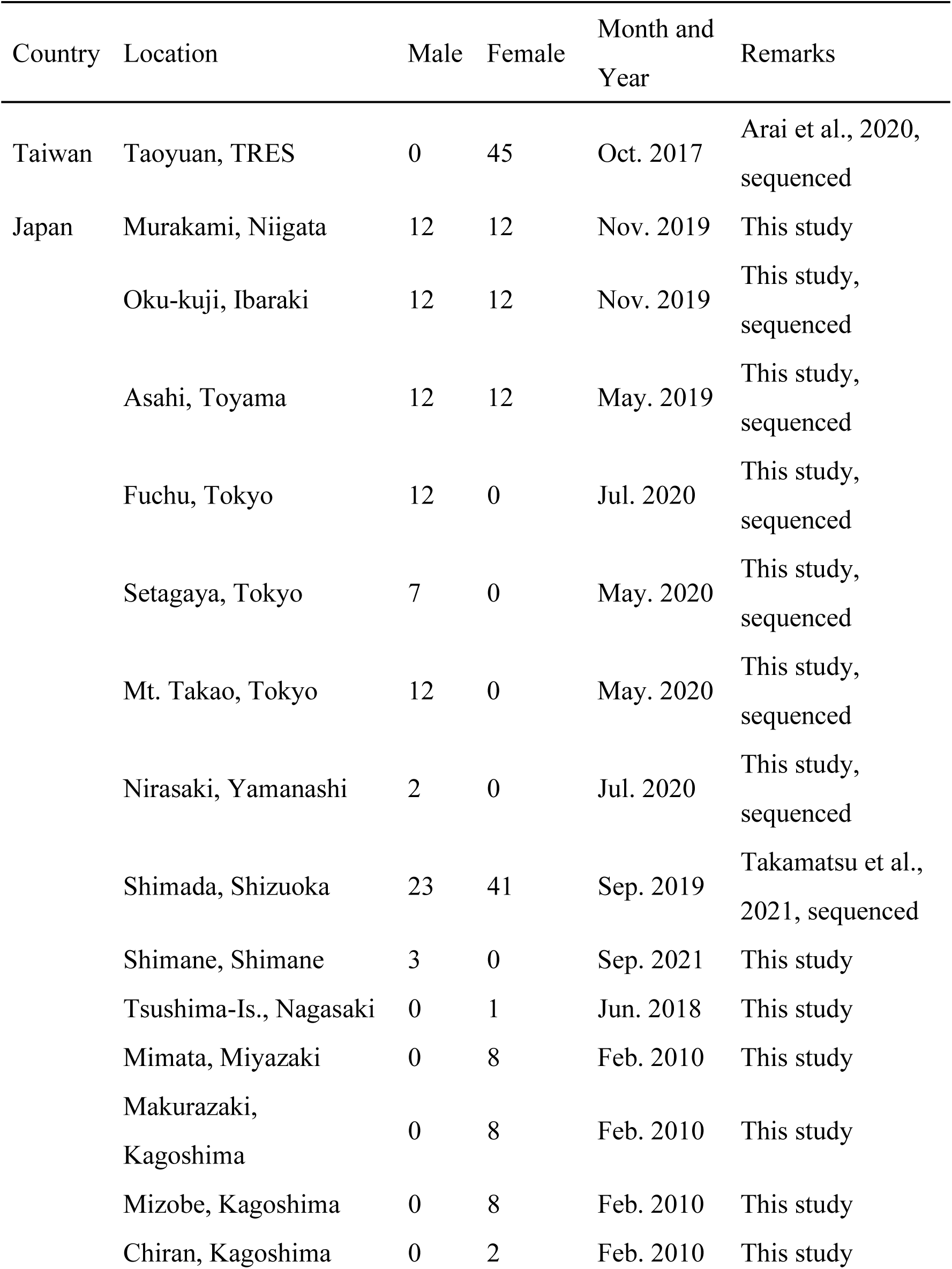
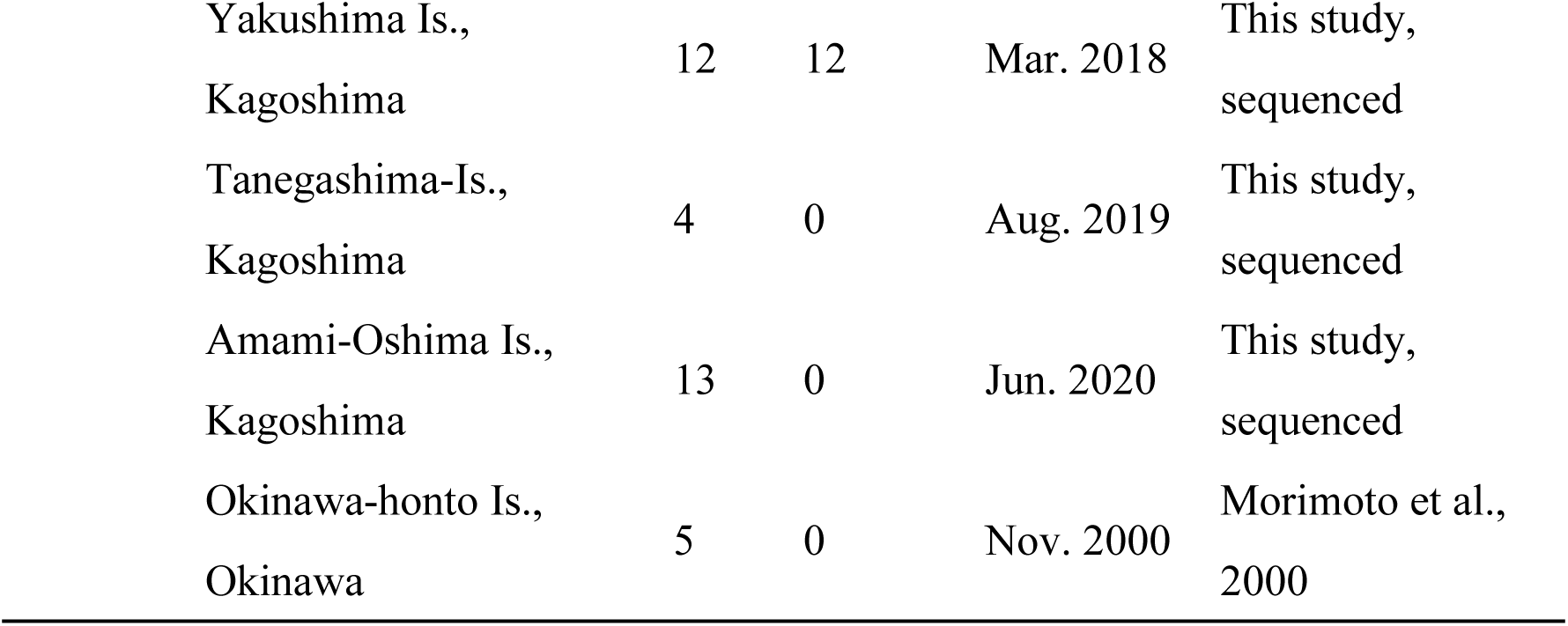
Sampling information of Taiwanese and Japanese *H. magnanima*

**Extended Data Table 3.**
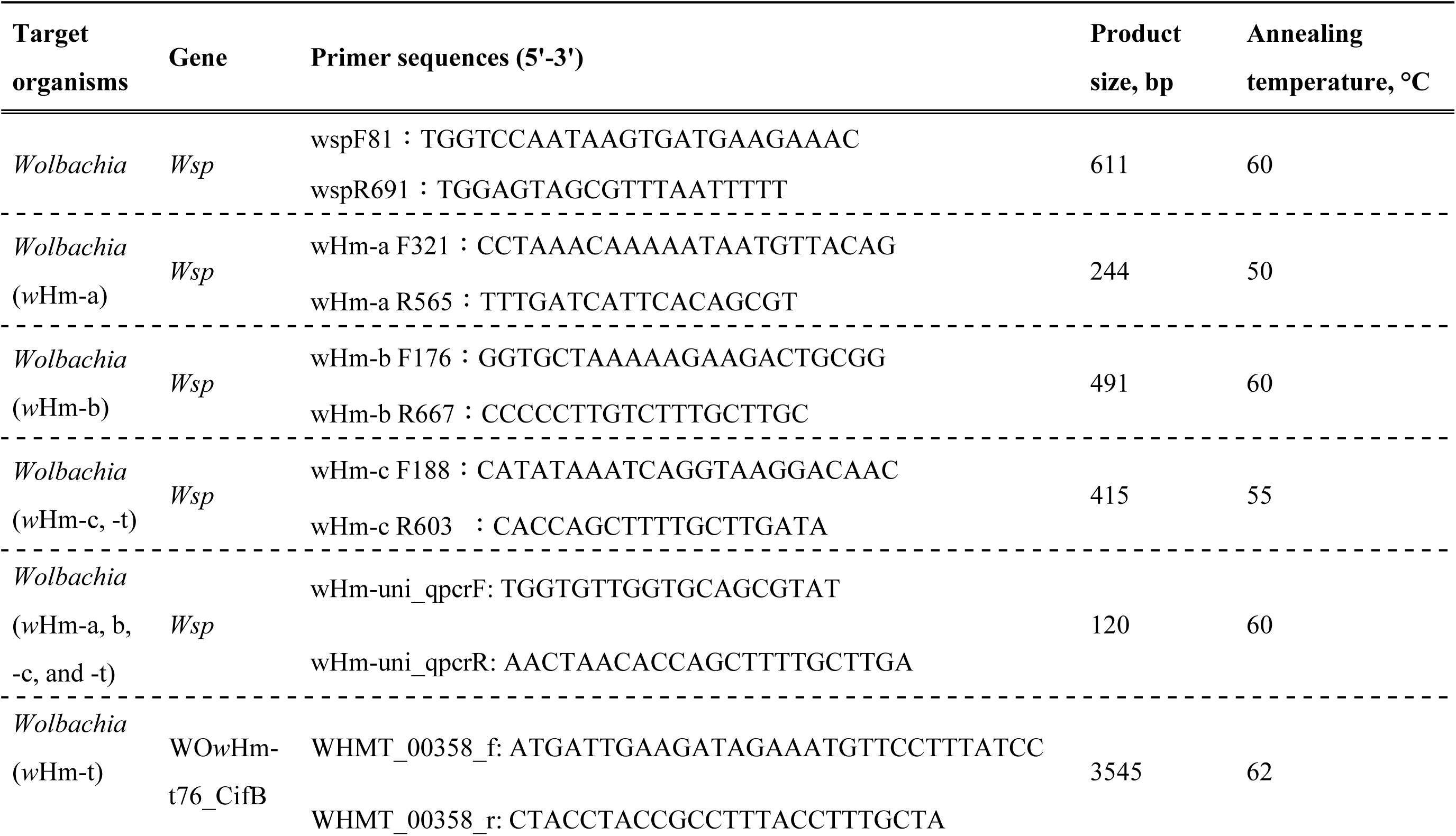
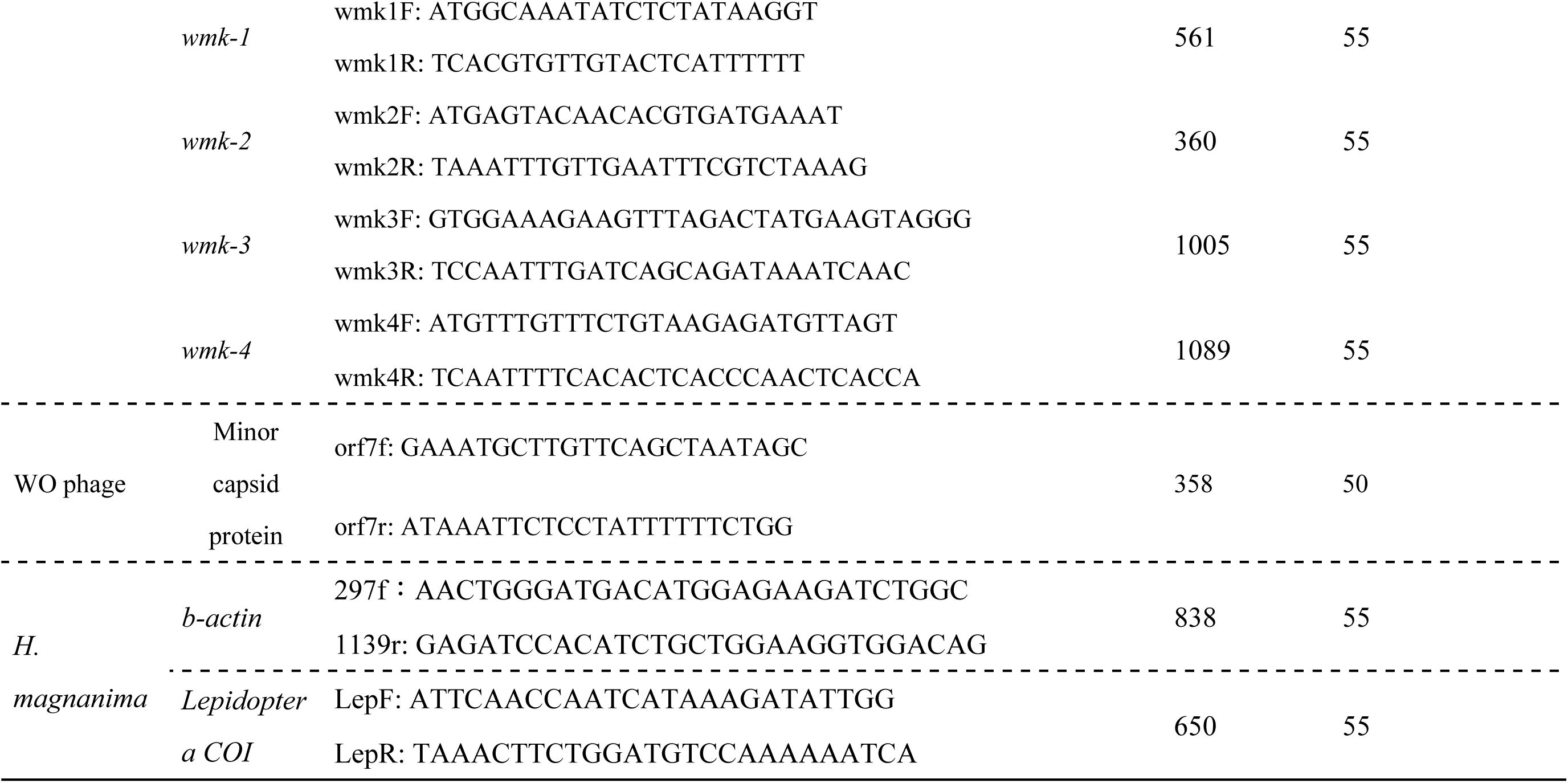
Primers for PCR detection.

## References

1. Sasaki, T. & Ishikawa, H. Production of essential amino acids from glutamate by mycetocyte symbionts of the pea aphid, *Acyrthosiphon pisum*. J. Insect Physiol. 41, 41–46 (1995).

2. Anbutsu, H. et al. Small genome symbiont underlies cuticle hardness in beetles. Proc. Natl Acad. Sci. U. S. A. 114, E8382–E8391 (2017).

3. Werren, J. H., Baldo, L. & Clark, M. E. *Wolbachia*: master manipulators of invertebrate biology. Nat. Rev. Microbiol. 6, 741–751 (2008).

4. Hurst, L. D. The incidences and evolution of cytoplasmic male killers. Proc. R. Soc. Lond. B 244, 91–99 (1991).

5. Hurst, G. D. D. et al. Invasion of one insect species, *Adalia bipunctata*, by two different male-killing bacteria. Insect Mol. Biol. 8, 133–139 (1999).

6. Andreadis, T. G. & Hall, D. W. Significance of transovarial infections of *Amblyospora sp*. (microspora: Thelohaniidae) in relation to parasite maintenance in the mosquito Culex salinarius. J. Invertebr. Pathol. 34, 152–157 (1979).

7. Fujita, R. et al. Late Male-Killing Viruses in Homona magnanima Identified as Osugoroshi Viruses, Novel Members of Partitiviridae. Front. Microbiol. 11, 620623 (2020).

8. Kageyama, D., Yoshimura, K., Sugimoto, T. N., Katoh, T. K. & Watada, M. Maternally transmitted non-bacterial male killer in *Drosophila biauraria*. Biol. Lett. 13, 20170476 (2017).

9. Majerus, M. E. N. & Hurst, G. D. D. Ladybirds as a model system for the study of male-killing symbionts. Entomophaga 42, 13–20 (1997).

10. Charlat, S. et al. Extraordinary flux in sex ratio. Science 317, 214–214 (2007).

11. Hayashi, M., Nomura, M. & Kageyama, D. Rapid comeback of males: evolution of male-killer suppression in a green lacewing population. Proc. R. Soc. Lond. B 285(1877), 20180369 (2018).

12. Werren, J. H. & Beukeboom, L. W. Sex determination, sex ratios, and genetic conflict. Annu. Rev. Ecol. Syst. 29, 233–261 (1998).

13. Zug, R. & Hammerstein, P. Still a host of hosts for *Wolbachia*: analysis of recent data suggests that 40% of terrestrial arthropod species are infected. PLOS ONE 7, e38544 (2012).

14. Ferri, E. et al. New insights into the evolution of *Wolbachia* infections in filarial nematodes inferred from a large range of screened species. PLOS ONE 6, e20843 (2011).

15. Zhou, W., Rousset, F. & O’Neil, S. Phylogeny and PCR–based classification of *Wolbachia* strains using *wsp* gene sequences. Proc. Biol. Sci. 265, 509–515 (1998).

16. Baldo, L. et al. Multilocus sequence typing system for the endosymbiont Wolbachia pipientis. Appl. Environ. Microbiol. 72, 7098–7110 (2006).

17. Jiggins, F. M., Hurst, G. D. D., Dolman, C. E. & Majerus, M. E. N. High-prevalence male-killing *Wolbachia* in the butterfly *Acraea encedana*. J. Evol. Biol. 13, 495–501 (2000).

18. Duplouy, A. et al. Draft genome sequence of the male-killing *Wolbachia* strain w Bol1 reveals recent horizontal gene transfers from diverse sources. BMC Genomics 14, 1–13 (2013).

19. Metcalf, J. A., Jo, M., Bordenstein, S. R., Jaenike, J. & Bordenstein, S. R. Recent genome reduction of *Wolbachia* in *Drosophila recens* targets phage WO and narrows candidates for reproductive parasitism. PeerJ 2, e529 (2014).

20. Arai, H., Lin, S. R., Nakai, M., Kunimi, Y. & Inoue, M. N. Closely Related Male-Killing and Nonmale-Killing *Wolbachia* Strains in the Oriental Tea Tortrix *Homona magnanima*. Microb. Ecol. 79, 1011–1020 (2020).

21. LePage, D. P. et al. Prophage WO genes recapitulate and enhance *Wolbachia*-induced cytoplasmic incompatibility. Nature 543, 243–247 (2017).

22. Beckmann, J. F., Ronau, J. A. & Hochstrasser, M. A *Wolbachia* deubiquitylating enzyme induces cytoplasmic incompatibility. Nat. Microbiol. 2, 17007 (2017).

23. Perlmutter, J. I. et al. The phage gene *wmk* is a candidate for male killing by a bacterial endosymbiont. PLOS Pathog. 15, e1007936 (2019).

24. Jaenike, J. Spontaneous emergence of a new *Wolbachia* phenotype. Evolution 61, 2244–2252 (2007).

25. Perlmutter, J. I., Meyers, J. E. & Bordenstein, S. R. A single synonymous nucleotide change impacts the male-killing phenotype of prophage WO gene wmk. eLife 10, e67686 (2021).

26. Harumoto, T. & Lemaitre, B. Male-killing toxin in a bacterial symbiont of *Drosophila*. Nature 557, 252–255 (2018).

27. Harumoto, T., Anbutsu, H., Lemaitre, B. & Fukatsu, T. Male-killing symbiont damages host’s dosage-compensated sex chromosome to induce embryonic apoptosis. Nat. Commun. 7, 12781 (2016).

28. Reynolds, L. A., Hornett, E. A., Jiggins, C. D. & Hurst, G. D. D. Suppression of *Wolbachia*-mediated male-killing in the butterfly *Hypolimnas bolina* involves a single genomic region. PeerJ 7, e7677 (2019).

29. Veneti, Z. et al. Loss of reproductive parasitism following transfer of male-killing *Wolbachia* to *Drosophila melanogaster* and *Drosophila simulans*. Heredity 109, 306–312 (2012).

30. Sasaki, T., Kubo, T. & Ishikawa, H. Interspecific transfer of *Wolbachia* between two lepidopteran insects expressing cytoplasmic incompatibility: a *Wolbachia* variant naturally infecting *Cadra cautella* causes male killing in *Ephestia kuehniella*. Genetics 162, 1313–1319 (2002).

31. Hornett, E. A. et al. Evolution of male-killer suppression in a natural population. PLOS Biol. 4, e283 (2006).

32. Arai, H. et al. Multiple infection and reproductive manipulations of *Wolbachia* in *Homona magnanima* (Lepidoptera: Tortricidae). Microb. Ecol. 77, 257–266 (2019).

33. Chrostek, E. & Teixeira, L. Mutualism breakdown by amplification of *Wolbachia* genes. PLOS Biol. 13, e1002065 (2015).

34. Masui, S. et al. Bacteriophage WO and virus-like particles in *Wolbachia*, an endosymbiont of arthropods. Biochem. Biophys. Res. Commun. 283, 1099–1104 (2001).

35. Fujii, Y., Kubo, T., Ishikawa, H. & Sasaki, T. Isolation and characterization of the bacteriophage WO from *Wolbachia*, an arthropod endosymbiont. Biochem. Biophys. Res. Commun. 317, 1183–1188 (2004).

36. Kent, B. N. & Bordenstein, S. R. Phage WO of *Wolbachia*: lambda of the endosymbiont world. Trends Microbiol. 18, 173–181 (2010).

37. Bordenstein, S. R. & Wernegreen, J. J. Bacteriophage flux in endosymbionts (*Wolbachia*): infection frequency, lateral transfer, and recombination rates. Mol. Biol. Evol. 21, 1981–1991 (2004).

38. Chafee, M. E., Funk, D. J., Harrison, R. G. & Bordenstein, S. R. Lateral phage transfer in obligate intracellular bacteria (*Wolbachia*): verification from natural populations. Mol. Biol. Evol. 27, 501–505 (2010).

39. Lindsey, A. R. I. et al. Evolutionary genetics of cytoplasmic incompatibility genes *cifA* and *cifB* in prophage WO of *Wolbachia*. Genome Biol. Evol. 10, 434–451 (2018).

40. Jones, J. D. & Dangl, J. L. The plant immune system. Nature 444, 323–329 (2006).

41. Hill, T., Unckless, R. L. & Perlmutter, J. I. Rapid evolution and horizontal gene transfer in the genome of a male-killing *Wolbachia*. bioRxiv (2020).

42. Hornett, E. A., Kageyama, D. & Hurst, G. D. D. Sex determination systems as the interface between male-killing bacteria and their hosts. Proc. Biol. Sci. 289, 20212781 (2022).

43. Hamilton, W. D. Extraordinary sex ratios. A sex-ratio theory for sex linkage and inbreeding has new implications in cytogenetics and entomology. Science 156, 477–488 (1967).

44. Harumoto, T., Fukatsu, T., & Lemaitre, B. Common and unique strategies of male killing evolved in two distinct *Drosophila* symbionts. Proc. Biol. Sci. 285, 20172167 (2018).

45. Sugimoto, T. N., & Ishikawa, Y. A male-killing *Wolbachia* carries a feminizing factor and is associated with degradation of the sex-determining system of its host. Biol. Lett. 8, 412–415 (2012).

46. Sugimoto, T. N., Kayukawa, T., Shinoda, T., Ishikawa, Y., & Tsuchida, T. Misdirection of dosage compensation underlies bidirectional sex-specific death in *Wolbachia*-infected *Ostrinia scapulalis*. Insect Biochem. Mol. Biol. 66, 72–76 (2015).

47. Fukui, T., Kawamoto, M., Shoji, K., Kiuchi, T., Sugano, S., Shimada, T., Suzuki, Y., & Katsuma, S. The endosymbiotic bacterium *Wolbachia* selectively kills male hosts by targeting the masculinizing gene. PLoS Pathog. 11, e1005048 (2015).

48. Arai, H. et al. Distinct Effects of the Male-Killing Bacteria Wolbachia and Spiroplasma and a Partiti-Like Virus in the Tea Pest Moth, Homona magnanima. bioRxiv (2022).

## Methods-only References

49. Iturbe-Ormaetxe, I., Woolfit, M., Rancès, E., Duplouy, A. & O’Neill, S. L. A simple protocol to obtain highly pure *Wolbachia* endosymbiont DNA for genome sequencing. J. Microbiol. Methods 84, 134–136 (2011).

50. Koren, S., Walenz, B.P., Berlin, K., Miller, J.R., Phillippy, A.M. Canu: scalable and accurate long-read assembly via adaptive k-mer weighting and repeat separation. Genome Res. 27, 722–736 (2017).

51. Li, H. Minimap2: pairwise alignment for nucleotide sequences. Bioinformatics 34, 3094–3100 (2018).

52. Bruce, J. W. et al. Pilon: An Integrated Tool for Comprehensive Microbial Variant Detection and Genome Assembly Improvement. PLoS ONE 9, e112963 (2014).

53. Chijiiwa, R. et al. Single-cell genomics of uncultured bacteria reveals dietary fiber responders in the mouse gut microbiota. Microbiome 8, 5 (2020).

54. Nishikawa, Y., et al. Massively parallel single-cell genome sequencing enables high-resolution analysis of soil and marine microbiome. bioRxiv (2020).

55. Tanizawa, Y., Fujisawa, T., & Nakamura, Y. DFAST: a flexible prokaryotic genome annotation pipeline for faster genome publication. Bioinformatics 34, 1037–1039 (2018).

56. Arndt, D., Grant, J. R., Marcu, A., Sajed, T., Pon, A., Liang, Y., & Wishart, D. S. PHASTER: a better, faster version of the PHAST phage search tool. Nucleic Acids Res. 44, W16–W21 (2016).

57. Kumar, S., Stecher, G., & Tamura, K. MEGA7: molecular evolutionary genetics analysis version 7.0 for bigger datasets. Mol. Biol. Evol. 33, 1870–1874 (2016).

58. Li, H. et al. The sequence alignment/map format and SAMtools. Bioinformatics 25, 2078–2079 (2009).

59. Sullivan, M. J., Petty, N. K., & Beatson, S. A. Easyfig: a genome comparison visualizer. Bioinformatics 27, 1 (2011).

60. Darling, A. C., Mau, B., Blattner, F. R., & Perna, N. T. Mauve: multiple alignment of conserved genomic sequence with rearrangements. Genome Res. 14, 1394–1403 (2004).

61. Arai, H., Ishitsubo, Y., Nakai, M., & Inoue, M. N. Mass-Rearing and Molecular Studies in Tortricidae Pest Insects. J. Vis. Exp. 181 (2022).

